# *PhyloVision*: Interactive Software for Integrated Analysis of Single-Cell Transcriptomic and Phylogenetic Data

**DOI:** 10.1101/2021.09.13.460142

**Authors:** Matthew G. Jones, Yanay Rosen, Nir Yosef

## Abstract

Recent advances in CRISPR-Cas9 engineering and single-cell assays have enabled the simultaneous measurement of single-cell transcriptomic and phylogenetic profiles. However, there are few computational tools enabling users to integrate and derive insight from a joint analysis of these two modalities. Here, we describe *PhyloVision:* an open source software for interactively exploring data from both modalities and for identifying and interpreting heritable gene modules whose concerted expression are associated with phylogenetic relationships. *PhyloVision* provides a feature-rich, interactive, and shareable web-based report for investigating these modules, while also supporting several other data and meta-data exploration capabilities. We demonstrate the utility of *PhyloVision* using a published dataset of metastatic lung adenocarcinoma cells, whose phylogeny was resolved using a CRISPR/Cas9-based lineage-tracing system. Together, we anticipate that *PhyloVision* and the methods it implements will be a useful resource for scalable and intuitive data exploration for any assay that simultaneously measures cell state and lineage.

## INTRODUCTION

Cellular lineages underlie several important biological phenomena - from embryogenesis, to differentiation, to cancer progression - and understanding the nature and dynamics of these lineages remains a central focus of research. Indeed, the piecing together of the developmental lineage of *C. elegans* by Sulston and colleagues via visual observations (Sulston *et al*., 1983) has facilitated decades of critical work using the deterministic development of *C. elegans* as a model system to study development (Packer, 2019), ageing (Kenyon, 2010), and even human diseases like neurodegeneration (Lu *et al*., 2014). Yet, many higher-order organisms cannot be studied by visual observation alone and thus a robust understanding of cell lineages underlying these organisms remains elusive. To this end several technologies have emerged to track cellular lineages over varying timescales, as reviewed in previous work (Kester and van Oudenaarden, 2018; McKenna and Gagnon, 2019).

Recently, the integration of CRISPR/Cas9-based engineering and single-cell sequencing has enabled the synthetic tracing of cellular lineages at unprecedented resolution (McKenna *et al*., 2016; Kalhor, Mali and Church, 2017; Raj *et al*., 2018; Chan *et al*., 2019; Bowling *et al*., 2020). Several of these technologies enable the simultaneous measurement of cell lineage and transcriptomic state via single-cell RNA-seq (scRNA-seq), thus creating exciting opportunities to study the transcriptional evolution of dynamic processes and motivating innovative approaches for integrating these two critical modalities (Wagner and Klein, 2020). As with high-dimensional measurements like those from scRNA-seq, it is clear that specialized, interactive tools for data exploration, visualization and analysis are necessary for realizing the full potential of these lineage tracing assays.

There exist several useful software tools for visualization of phylogenetic or lineage tracing data. For example, the interactive Tree of Life (iTOL; (Letunic and Bork, 2021)) is a scalable web-server-based tool that allows users to upload tree structures and various annotation files for interactive viewing. Yet, to indefinitely host and share these reports requires a paid subscription. More recently, CeLaVi was introduced as a publicly available software tool for generating interactive, web-based reports expressly for cell-lineage viewing (Salvador-Martínez *et al*., 2021). While both tools are scalable to up to thousands of cells and versatile for integrating various data modalities (e.g., gene expression measurements and spatial location) with phylogenies for visualization, they do not offer capabilities for joint analysis and automated interpretation of information on lineages and gene expression.

Here, we introduce *PhyloVision:* an open-source, interactive analysis and visualization tool that is expressly built for integrating single-cell gene expression and lineage data. *PhyloVision* builds on useful existing work, like iTOL and CeLaVi, for interactively visualizing phylogenies while possibly overlaying the expression of individual genes. In addition to this, however, *PhyloVision* also employs other analysis frameworks developed by our group for automated interpretation of the variation in gene expression, provided the lineage structure. Specifically, *PhyloVision* supports features that identify heritable gene expression programs and interpret these programs using gene signature enrichment analysis.

To demonstrate the utility of *PhyloVision*, we apply it to a clone of 1,127 cells from a recent CRISPR/Cas9 lineage-tracing dataset investigating metastatic spread in a mouse model of lung adenocarcinoma. In doing so, we show that the derived statistics and web-based user interface can be used to effectively characterize subpopulations within this aggressive tumor population. These molecular characterizations, not discussed in the original study, can be used to generate hypotheses about how metastatic ability evolves within a tumor subpopulation.

*PhyloVision* is distributed publicly on Github at https://github.com/YosefLab/VISION. Along with the software, we include several tutorials and example reports of published datasets allowing users to explore the user-interface. Additionally, we include a detailed manual and description of the user-interface in the Supplementary Materials.

## RESULTS

### *PhyloVision* is an integrated pipeline for interactive analysis of single-cell expression and lineage profiles

*PhyloVision* is simultaneously a tool for interactive exploration of multimodal single-cell lineage tracing data using our web-based front end and for analysis of the evolutionary dynamics of expression data. Our interactive web-based report is built on our *VISION* front-end (DeTomaso *et al*., 2019). Here, we have developed an interactive phylogeny viewer and have integrated it into the default interface, enabling the user to perform various manipulations on the observed tree (e.g., node collapsing) and to select groups of nodes (cells) for viewing on a low-dimensional embedding of the respective scRNA-seq data or for differential expression analysis (Fig 1). The dynamic phylogeny viewer is scalable, allowing low-latency selections and sub-tree collapsing for large trees (we tested trees of up to ∼4,000 leaves). Importantly, a web-report generated by a user can be viewed locally, shared privately amongst colleagues, or staged publicly on a web server.

**Figure 1:**
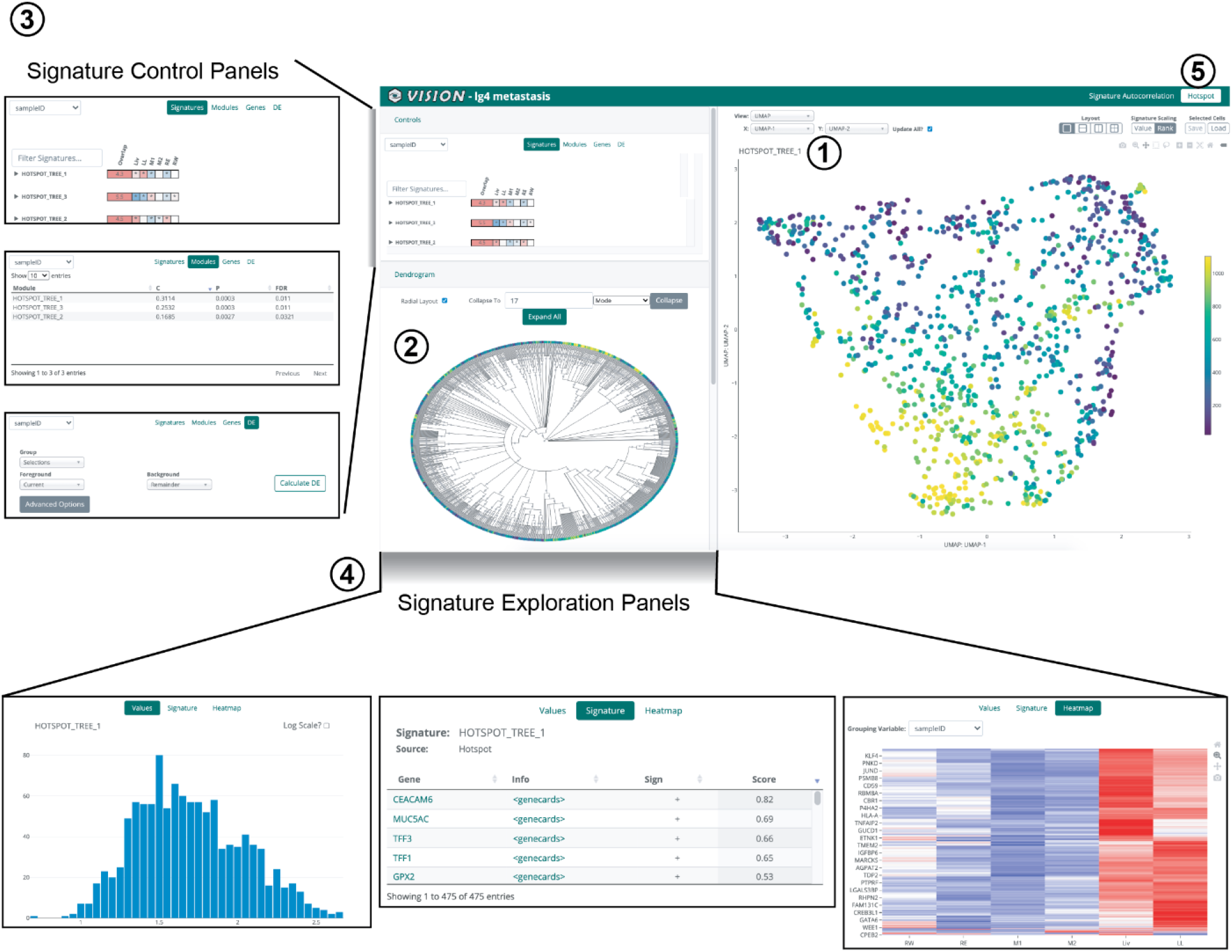
Overview of the *PhyloVision* interactive UI. *PhyloVision*’s user-interface (UI) is web-based, feature-rich report that can be hosted locally or externally. *PhyloVision* incorporates four main panels into viewing: first, a panel for visualization of two-dimensional single-cell RNA-seq projections or coordinates (e.g., from spatial transcriptomics datasets; inset 1). Second, a panel for interactive visualization of a phylogeny relating all cells (inset 2) that enables selection, collapsing, and variable layouts (radial or linear). Third, a control panel for selecting values to be overlaid onto the phylogeny and two-dimensional visualization panel, evaluating statistics associated with each signature or module, plotting a gene’s expression, or performing differential expression analysis (inset 3). Note that here the “*Hotspot”* mode is shown, which operates “bottom up”: first finding heritable gene modules and then analyzing their over-representation (enrichment) in user-provided signatures. The alternative, “Vision”, mode that operates directly on the user-signatures is described in (DeTomaso *et al*., 2019). Users can control the analysis mode by toggling between “Signature Autocorrelation” and “Hotspot” (inset 5). Fourth, an exploration panel for inspecting the value distribution, gene membership, and expression heatmaps of each user-provided gene signature or automatically-identified Hotspot module (inset 4).

While the interactive web-report is a useful tool for data exploration, the *PhyloVision* pipeline additionally supports statistical analysis for deriving joint insight from the expression and lineage data (Fig S1). In this, *PhyloVision* takes as input (at a minimum) an expression data matrix, a phylogeny, and a collection of gene signatures, each representing a certain pathway or a transcriptional response to a certain change in conditions (as publicly available from resources like MSigDB (Subramanian *et al*., 2005)). In one mode of analysis, *PhyloVision* will conduct a phylogenetic autocorrelation analysis with the user-defined gene signatures. Here, for a given signature, a score will be computed for each cell with *VISION* (*DeTomaso et al*., *2019*), indicating the cumulative activity of the respective genes (note that the score also accounts for “signed” transcriptional response signatures, in which one subset of genes is marked as up-regulated and another as down-regulated). An autocorrelation statistic will be then computed by evaluating the consistency of the signature scores between nearby cells on the phylogeny (using the Geary’s C statistic (Geary, 1954) as in (DeTomaso *et al*., 2019)). With this analysis, a user can identify gene signatures that are significantly associated with the tree structure, suggesting evolutionary patterns of interest. Our implementation of *VISION* also adds capabilities to visualize and analyze user-provided meta-data. This mode allows the user to identify cell-level properties (e.g., tissue of origin, or the extent of somatic mutations) whose variation is consistent with the tree structure, and which might therefore represent sub-clonal, heritable phenotypes.

In another mode of analysis, users can identify *de novo* gene modules (i.e., not guided by pre-defined signatures) that are learned from the intersection of phylogeny and expression data using our *Hotspot* algorithm (DeTomaso and Yosef, 2021). Briefly, this analysis uses autocorrelation to identify individual genes whose expression is consistent with the phylogeny - namely genes that are expressed at a more similar level in phylogenetically adjacent cells than in cells that are distant. It then uses a pairwise extension of the autocorrelation statistic to arrange these genes into modules whose expression patterns on the phylogeny are similar, thus representing conserved transcriptomic modules that each operate in a concerted fashion. The *PhyloVision* pipeline adds interpretability to the *Hotspot* modules by assessing the overlap between their respective gene sets and the user-provided gene signatures. *PhyloVision* provides a quantification of this overlap with an enrichment statistic and an assessment of statistical significance. Together, these two analyses enable a user to identify important sources of transcriptomic variation on the phylogeny, as well as discover new and interpretable gene sets.

### Case study: Analysis of a metastatic lung adenocarcinoma tumor with *PhyloVision*

To demonstrate *PhyloVision*’s usefulness in interrogating data from multimodal single-cell lineage tracing technologies, we applied the pipeline to a clone from our recently published dataset in which we studied the metastatic behavior of an aggressive human lung adenocarcinoma cell line in a xenograft mouse model (Quinn *et al*., 2021).

As previously described, we used our CRISPR/Cas9 lineage tracing technology (Chan *et al*., 2019) to trace approximately 100 clones over the course of 2.5 months as each clone metastasized between tissues in a mouse model of lung adenocarcinoma. In this analysis, we used the reconstructed single-cell phylogenies from the original study, which were inferred with the Cassiopeia package (Jones *et al*., 2020). In these phylogenies, each leaf corresponds to a cell with data corresponding to the single-cell expression profile and the tissue from which it was sampled. In the original study, we described how metastatic rates of single cells could be inferred directly from these phylogenies and combined with expression profiles to identify transcriptional regulators of this process.

In the present analysis, we evaluated a clone of 1,127 cells using the *PhyloVision* pipeline (interactive report available at http://s133.cs.berkeley.edu:8224/Results.html). In this case study, we used signatures downloaded from MSigDB (Subramanian *et al*., 2005), and focused on the results from the *Hotspot* analysis. *Hotspot* identifies three non-overlapping modules of genes (Fig 2A). To evaluate the cumulative activity of each module at each cell in the dataset, we computed module scores for every cell using the signature-scoring procedure in *VISION* (see Methods). Interestingly, when compared to the metastatic rate inferred from the phylogeny (which is provided as meta-data to this session), we observe that one module is negatively correlated (Module 1; Pearson’s *ϱ =* -0.30, Fig 2B left) while one is positively correlated with this metastatic rate (Module 2; Pearson’s *ϱ =* 0.26, Fig 2B middle); Module 3 does not correlate with the metastatic rate in either direction (Pearson’s *ϱ* = 0.07, Fig 2B right). These results therefore point to candidate transcriptional programs (each represented by a module) that are heritable and are associated with different metastatic abilities.

**Figure 2:**
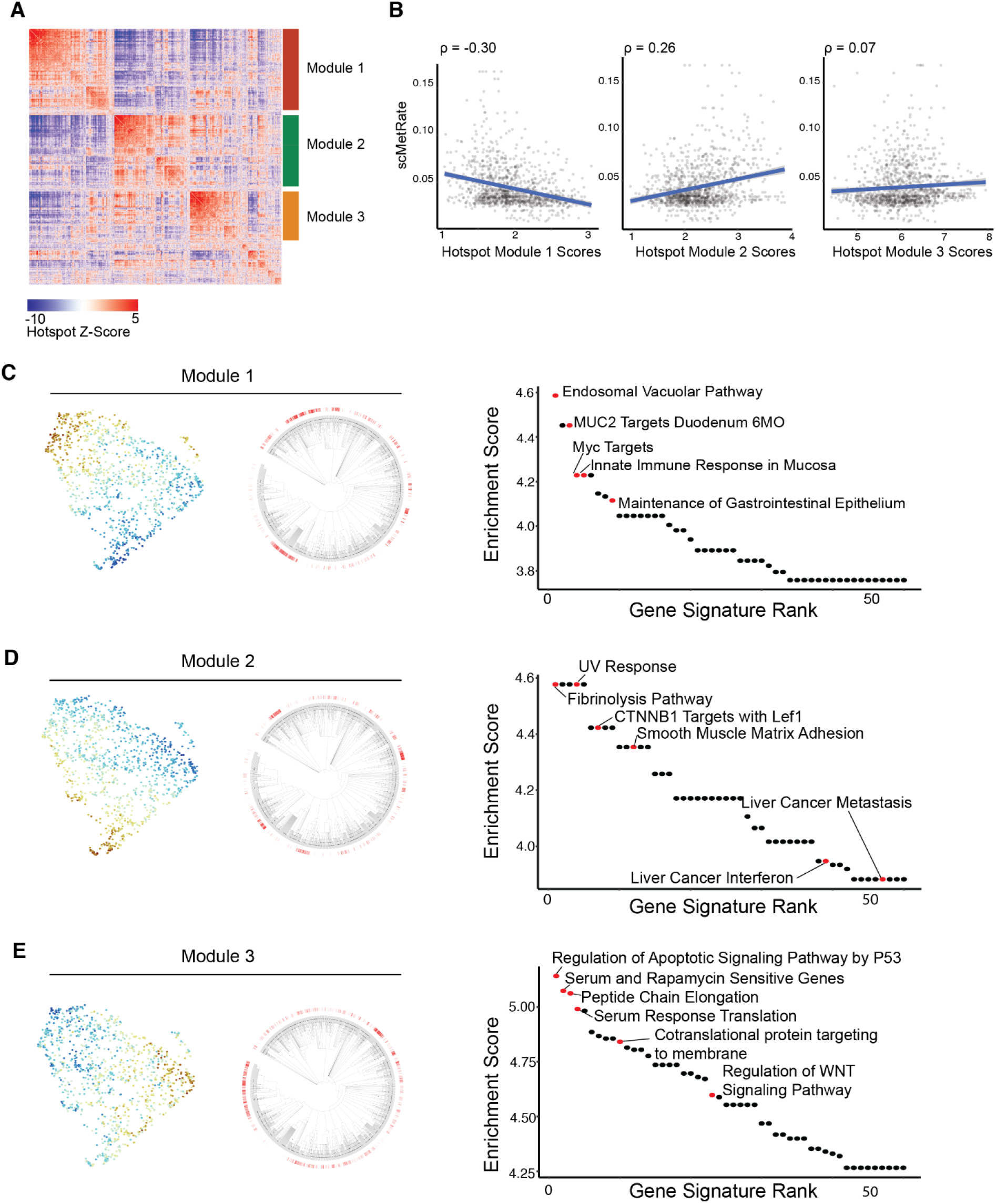
*PhyloVision* analysis identifies gene modules associated with metastasis and tumor progression. A. Heatmap of Z-normalized pairwise autocorrelations between *Hotspot*-selected genes, clustered into three non-overlapping modules (genes not grouped into a *Hotspot* gene module do not have a bar annotation). Note that pairwise autocorrelations can be interpreted as a smooth and more robust estimation of a standard pairwise correlation (DeTomaso and Yosef, 2021). B. Hotspot module scores and the single-cell metastasis rate (scMetRate) are plotted against one another. Pearson’s correlation coefficients are indicated above each scatterplot. C–E. Interpretation of *Hotspot* gene modules for Module 1 (C), Module 2 (D), and Module 3 (E). Module scores are overlaid onto the UMAP of scRNA-seq data and phylogenies. The enrichment score between gene signature and module gene set (defined as the ratio between their observed and expected overlap) is shown for the top 50 signatures. Selected gene signatures are highlighted and annotated.

To interpret the biological signal intrinsic to these *Hotspot* modules, we first projected the single-cell transcriptomic profiles onto two dimensions using Uniform Manifold Approximation and Projection (UMAP; (McInnes, Healy and Melville, 2018)) and overlaid the *Hotspot* modules scores (Fig 2C-E). Upon inspection, we observed that the *Hotspot* modules localized to distinct regions of the two-dimensional projection, and therefore marked different cellular states. We observed a similar pattern, on the phylogenies themselves, with each Hotspot module marking a specific set of sub-clades.

Next, we examined which gene signatures had significant overlap with each module. In the module negatively correlated with metastatic rate (Module 1), we found significant enrichments corresponding to the innate immune response, maintenance of the gastrointestinal epithelium, and the endosomal vacuolar pathway, amongst others (all FDR < 1e-3, hypergeometric test; Fig 2C). Together, these gene sets indicate that less metastatic cells in this clone are characterized by antigen processing and presentation and maintenance of an epithelial-like state, supporting the hypothesis that tumor progression is required for metastatic competency in lung adenocarcinoma (Caswell *et al*., 2014).

On the other hand, the module positively correlated with metastatic rate (Module 2) had significant enrichments in gene signatures associated with fibrinolysis, UV response, smooth muscle adhesion, inflammatory response, and other metastatic signatures (e.g., liver cancer metastasis; all FDR < 0.05, hypergeometric test; Fig 2D). These gene signatures therefore point to several mechanisms of enhancing metastatic rates in this clone, such as fibrinolysis and up-regulation of adhesion molecules. The diversity of these signatures underscore the importance of inflammatory signaling, ECM remodelling, and cell-adhesion discussed in our previous work (Quinn *et al*., 2021).

Module 3 was characterized by a set of gene signatures associated with tumor progression but distinct from the metastatic rate. Intriguingly, the gene sets that significantly overlapped with Module 3 included those that regulated apoptosis and WNT-signaling, as well as those indicating sensitivity to serum and rapamycin treatment (all FDR < 1e-3, hypergeometric test; Fig 2E). These observations suggest that Module 3 distinguishes cells in an altogether different metabolic state uncorrelated with metastatic ability but appearing to be associated with increased survival ability and cell proliferation, perhaps due to relatively high amounts of *KRAS* signaling. These results also support the finding that combination therapies involving inhibitors of mTOR, which sits downstream of *RAS* signaling, is a viable therapeutic opportunity in *KRAS-*driven non-small-cell lung cancer (Vasan, Boyer and Herbst, 2014). Future work may investigate if this therapy-sensitive population is mainly characterized by differential metabolic signaling, perhaps leveraging recent work modeling metabolic profiles from scRNA-seq data (Wagner *et al*., 2021).

Overall, these results indicate that the *PhyloVision* joint analysis of single-cell expression and lineage provides an efficient approach for dissecting phylogeny-based transcriptional heterogeneity. In the case-study above, we demonstrate that the analysis pipeline and web-based UI were able to identify gene programs associated with increased or decreased metastatic ability, and altogether new programs not previously described in the original study. As a whole, this analysis provides testable hypotheses and intricate molecular characterizations for subpopulations in a single clone.

## DISCUSSION

We introduced *PhyloVision*, a new tool for the integrated analysis of scRNA-seq and single-cell lineage tracing data. *PhyloVision* offers a feature-rich, user-friendly, and interactive web report for exploring the evolutionary underpinnings of scRNA-seq profiles. Moreover, *PhyloVision* is embedded within useful analysis pipelines, thus enabling rapid characterization of interesting structure in the high-dimensional data. In this way, *PhyloVision* is the first interactive tool of its kind to provide a bridge between single-cell analysis tools and single-cell lineage tracing data. We show the effectiveness of this approach in a case study of a single clone from our recent work, illustrating rich heterogeneity in this dataset previously unexplored.

In addition to the Hotspot analysis presented here, *PhyloVision* includes other analysis features such as identification of heritable meta-data (e.g., metastatic rate) and its transcriptional correlates (using gene signatures and hotspot modules), identification of differentially expressed genes (e.g., by manually choosing sub-clones to be compared), and visualization of gene expression and meta data while stratifying the cells according to the phylogeny (e.g., with histograms corresponding to different clades).

While the case study presented in this work focused on CRISPR/Cas9-based lineage tracing, we expect that this tool will also be useful for any type of data that jointly measure cell lineage and transcriptomic state such as those from B cell phylogenies (inferred from their antigen receptor) or single-cell phylogenies built from whole-genome sequencing of tumor samples. Moreover, the shareability of generated web-based reports allow collaborators without computational experience to smoothly explore high-dimensional datasets. Taken together, we anticipate that *PhyloVision* will be useful across applications and technologies and will provide critical support for the interpretation of these multimodal datasets.

## AVAILABILITY AND IMPLEMENTATION

*PhyloVision* is publicly available under the MIT license at https://github.com/YosefLab/Vision. The Plotly-enabled javascript phylogeny plotting and viewing library is publicly available at https://github.com/YosefLab/PhyloPlot. Example reports are available at http://s133.cs.berkeley.edu:8224/Results.html (metastasis xenograft dataset from (Quinn *et al*., 2021)) and http://s133.cs.berkeley.edu:8225/Results.html (mouse embryogenesis from (Chan *et al*., 2019)).

## ACKNOWLEDGEMENTS

The authors would like to thank Alex Khodaverdian, Richard Zhang, Sebastian Prillo, Joseph Min, David DeTomaso and the members of the Yosef lab for their helpful discussions in the development of this project. The authors would also like to thank Galen Xing, Liron Grossman, and Shirley Hui for helpful testing and feedback of the software. This work was supported by UCSF Discovery Fellowship (M.G.J.) and NIH-NIAID U19AI090023 (N.Y.). N.Y. is also a member of the Chan Zuckerberg Biohub investigator program.

## AUTHOR CONTRIBUTIONS

All authors contributed to the design of software and analysis. N.Y and M.G.J. conceived of this project, and N.Y. directed the work. Y.R. and M.G.J. wrote the software. M.G.J. performed the analysis on the xenograft cancer dataset. All authors wrote and approved the final manuscript.

## DECLARATION OF INTERESTS

N.Y. is an adviser and/or has equity in Cellarity, Celsius Therapeutics, and Rheos Medicines. All other authors declare no competing interests.

## METHODS

### The PhyloVision pipeline

*PhyloVision* builds on the *VISION* analysis toolkit for signature autocorrelation analysis (DeTomaso *et al*., 2019). As input, *PhyloVision* requires a gene expression matrix (typically count-normalized, but not log-normalized), a set of signature gene sets (e.g., publicly available from sources like MSigDB), and a phylogeny (stored as a tree structure in the *ape* R package (Paradis and Schliep, 2019)). Amongst other data that can be optionally passed into *PhyloVision* are numerical and categorical metadata, as well as a two-dimensional projections of the cells for visualization purposes (e.g. from t-distributed stochastic neighbor embedding [tSNE] (van der Maaten, 2008) of the main principal components or of a latent representation learned by methods such as scVI (Lopez *et al*., 2018)). A related optional input is a low-dimensional (usually more than 2 dimensions) representation of the data (e.g., top principal components or the scVI latent space), which is used for clustering the cells (to see how cell clusters are utilized, see section “Analysis of metadata”) and evaluating the consistency of signature scores. If the latter inputs are not provided, then PhyloVision computes the low-dimensional representation using PCA and the further 2-dimensional projections using tSNE and UMAP (though other algorithms are supported as well, which users can specify). Finally, users can also specify additional pre-computed Hotspot objects that can be used for the Hotspot analysis (as described below).

The signature-centric (VISION) part of the analysis pipeline in *PhyloVision* (invoked by default when running *PhyloVision*) begins by clustering cells according to substructure on the tree (see below, section “Generating stratification (clustering) of the cells according to the phylogeny”), computing scores for each user-provided signature and for every cell (computed using cell-level Z-scores as in (DeTomaso *et al*., 2019)). It then assesses the agreement between the signature scores and the phylogeny with a phylogenetic autocorrelation statistic (see section entitled “Phylogenetic autocorrelation analysis”). Signatures that are significantly auto-correlated (i.e., cells that are nearby in the phylogeny have more similar scores than expected by chance) are clustered using the previously described *VISION* pipeline (DeTomaso *et al*., 2019) and displayed in the web-based report in a collapsable table for each cluster (Figure 1, inset 3). The user can explore the scores of these gene signatures by overlaying their scores on the two-dimensional representation of the gene expression data (Figure 1 inset 1) and the phylogeny (Figure 1 inset 2), viewing histograms of these scores possibly subsetted according to the phylogeny (Figure 1 inset 4), and browsing through the respective genes (Figure 1 panel 4; where genes are ranked their covariance with the signature score).

*PhyloVision* additionally supports a phylogenetic analysis with *Hotspot*, which can be invoked with the *runHotspot* function. This function interfaces with the *Hotspot* tool, implemented in Python, using the *reticulate* R package. Using the previously described *Hotspot* pipeline (DeTomaso and Yosef, 2021), *PhyloVision* first identifies individual genes whose expression is coherent with the tree structure (i.e., nearby cells in the tree express the gene at a similar level, as compared to random chance). It then performs a two-dimensional autocorrelation analysis to group the identified genes into modules. Finally, *PhyloVision* computes an enrichment statistic between user-defined gene signatures and the identified *Hotspot* modules (see section entitled “Assessment of statistical significance of *Hotspot* gene module and gene signature overlap”). Gene signatures with significant overlap are included in the web-based report in a collapsable table for each *Hotspot* module (Figure 1 inset 3). The user can explore the scores of these gene signatures or the score of entire hotspot modules by overlaying their cumulative expression (computed identically to user-specified signatures, as in (DeTomaso *et al*., 2019)) on the 2-dimensional representation of the gene expression data (Figure 1 inset 1) and the phylogeny (Figure 1 inset 2), viewing histograms of these scores possibly subsetted according to the phylogeny (Figure 1 inset 4), and browsing through the respective genes (Figure 1 panel 4; where genes are ranked by covariance as with user-specified signatures).

### Phylogenetic autocorrelation analysis for gene sets (signature-centric analysis)

To compute the extent to which a value (e.g., signature score, module score, or continuous covariate) can explain the cellular relationships on the phylogeny, we make use of the Geary’s *C* statistic for local autocorrelation. This statistic is defined as

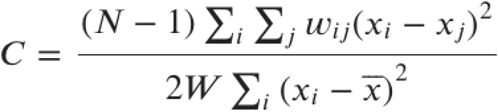

where *w*_*ij*_ represents the cophenetic distance (i.e., distance between cells using the branch lengths of the user-specified phylogeny) between cells *i* and *j, x*_*i*_ is a value of interest, *N* is the total number of cells, and *W* is the sum of all weights. (A small amount of random noise is introduced to the cophenetic distances to break ties.) In our case, the value of interest (i.e. *x*) are the ranks of the normalized signature score in each cell, as defined previously (DeTomaso *et al*., 2019). We report *C’ = 1 - C* such that a score of 1 indicates perfect autocorrelation and 0 means no autocorrelation. While the *C’* statistic provides an effect size, we evaluate the significance of gene signature scores with an empirical p-value (FDR corrected with the Benjamini-Hochberg procedure (Benjamini and Hochberg, 1995)), comparing the signature score to a background of randomly generated signatures as described in our previous work (DeTomaso *et al*., 2019). In the collapsable tables displayed on the UI (Figure 1, inset 3), we retain only gene signatures with an FDR < 0.05 and report both C’ statistics and the FDR.

In the phylogenetic autocorrelation analysis, we utilize a K-nearest neighbor (KNN) graph where weights are only non-zero between a cell *i* and it’s closest *k* neighbors. Specifically, if cell *j* is not a k-nearest neighbor of cell *i*, then *w*_*ij*_ is taken to be 0. K-nearest neighbors of cell *i* are found using the distances on the phylogeny, where the distance between cells *i* and *j* is defined as the sum of the edge lengths on the path between the two cells. If edge lengths for the dendrogram are not provided, every edge is defaulted to length 1. Ties are randomly broken.

### Phylogenetic autocorrelation analysis for individual genes and the identification of modules (HotSpot analysis)

*PhyloVision* employs a gene-level clustering into modules using the *Hotspot* autocorrelation analysis (DeTomaso and Yosef, 2021) on the user-defined phylogeny. (Here we describe how the algorithm is applied in the *PhyloVision* pipeline, for mathematical details please refer to our previous work (DeTomaso and Yosef, 2021).) First, using the cophenetic distances on the tree (defined as the phylogenetic distance separating cells), a *K-*nearest neighbor (KNN) graph is constructed (using a default *K = sqrt(N)*, where *N* is the number of cells). Then, genes are selected that are significantly autocorrelated with the phylogenetic KNN graph using the “*compute_autocorrelations*” function in *Hotspot*. By default, the depth-adjusted negative binomial (“danb”) model is used and the top 1000 genes are selected (as measured by *Hotspot’s* Z-transformed Geary’s C) that pass an 0.05 FDR threshold, though both these parameters can be controlled by the user. Then, genes are grouped into modules by clustering the pairwise autocorrelation matrix computed with *Hotspot’s “compute_local_correlations”* function. By default, we use a minimum gene threshold of 20 and a clustering FDR of 0.5, though both parameters are controllable by the user. Gene signatures corresponding to the genes in a module are added to the *PhyloVision* object. Additionally, for each module and user-specified gene-signature pair, a new signature is created by computing the gene overlap and added to the *PhyloVision* object.

### Assessment of statistical significance of the overlap between Hotspot modules and user-provided gene signatures

Given a full set of *N* genes, *PhyloVision*’s compares the genes in module set *M* identified by *Hotspot* and the genes in an existing signature set *S* by first computing an enrichment statistic:

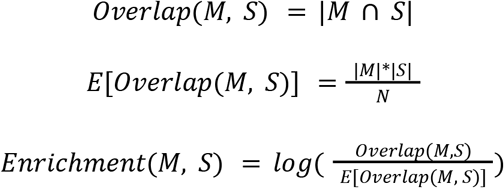

Then, we assess significance using a hypergeometric test with *R*:

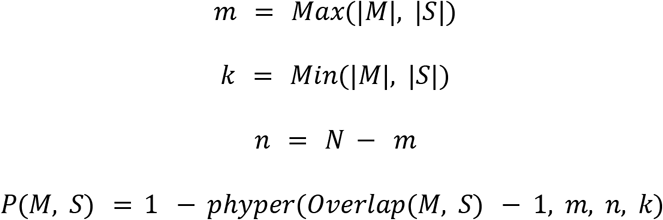

Where *P(M,S)* indicates the p-value of the overlap. P-values are then FDR-corrected using the Benjamini-Hochberg procedure (Benjamini and Hochberg, 1995).

### Analysis of metadata

*PhyloVision* utilizes the tools in *VISION* to conduct autocorrelation analysis on metadata. In this, *PhyloVision* analyzes metadata differently depending on if is numerical (i.e., continuous covariates like the number of genes detected in a cell) or categorical (i.e., discrete covariates like the batch in which a cell profile was sampled). While most metadata is specified by the user, *PhyloVision* additionally clusters the cells into subclades using a tree-based clustering method (see section below entitled “Generating stratification (clustering) of the cells according to the phylogeny”).

In the context of numerical metadata, *PhyloVision* utilizes an approach to assess autocorrelation and significance identical to that of gene-signatures (see above, “Phylogenetic autocorrelation analysis for gene sets (signature-centric analysis), with the exception that scores need to be computed as they are provided as precomputed scores by the user.

In the context of categorical data, *PhyloVision* uses the Cramer’s *V* statistic as an autocorrelation statistic. As described in our previous work (DeTomaso *et al*., 2019), the Cramer’s *V* statistic is a transformation of a chi-squared test statistic, computed on the local neighborhood of each cell. Specifically, for each cell *i*, we compute a local proportion of each variable value *m* across its *K* phylogenetic neighbors (indexed by *j*) :

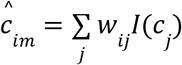

where *w*_*ij*_ are computed as above in the gene-centric analysis (see section “Phylogenetic autocorrelation analysis for gene sets (signature-centric analysis)”), *c*_*j*_ represents the value of the discrete variable of interest in cell *j* and *I*_*m*_*(x)* is an indicator function that takes on the value of 1 if *x == m* and 0 otherwise. From these values, a contingency table ***X*** is computed as

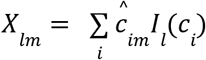

The chi-squared test is then performed on this contingency table ***X*** to estimate a *p-*value. From the chi-squared test statistic, *t*, the Cramer’s *V* statistic is computed as

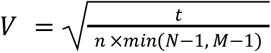

where *n* is the sum of all the values in the contingency table ***X***, *N* is the number of rows in the contingency table and *M* is the number of columns.

### Visualization of phylogenies

*PhyloVision*’s web-based user interface displays the provided phylogeny using a custom Plotly Javascript package (PhyloPlot.js) in either a radial or linear layout. Users are able to select individual cells and clades on the phylogeny for use in Differential Expression and to view on the UMAP. Leaves reflect the same cell-coloring as the UMAP. Users are able to collapse clades to summary nodes, using the mode, arithmetic mean, geometric mean or median of the numerical values selected for the node’s leaves. Users can also collapse nodes by depth from the root of the tree. The phylogeny in both radial and linear layouts is converted to ultrametric edge lengths using the following formula:

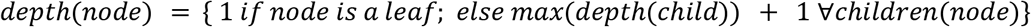

### Generating stratification (clustering) of the cells according to the phylogeny

In many cases, a phylogeny can be used to stratify groups of cells into specific groups by “cutting” the tree at a specific depth and assigning cells to subclonal lineages. For example, if a phylogeny describes a differentiation process between several cell types, a clustering of cells into subclades might yield cell-type specific clusters. *PhyloVision* performs not only a clustering of the phylogeny but also an assessment of how meaningful the clustering is on the latent space.

*PhyloVision* clusters cells on a phylogeny by performing a breadth-first search over internal nodes. Specifically, the algorithm maintains a queue of internal nodes and updates the queue by popping off the internal node with the largest child clade size, and adding it’s immediate children to the queue. This algorithm begins with the root node, and terminates once the queue has reached a target length (defaulted to 10) at which point *PhyloVision* merges the smallest clade with its neighbor until the exact target number is reached. The algorithmic pseudocode is detailed as follows:

~~~
**Cluster-Phylogeny** (target := 10)
    Queue := Priority Queue {Node : Maximum size of Node’s child clades}
    Insert the root into the queue
    While length(queue) <= target do:
      Node := queue.pop(1)
      Insert Node’s children to Queue
    Clusters := {[Children]} ∀Nodes ∈Queue
    While length(Clusters) > target do:
      Select smallest cluster from Clusters
      Merge that cluster with its phylogenetically nearest cluster
    Return Clusters
~~~

### Differential expression analysis

Differential Expression is performed using a Wilcoxon Rank Sum Test on the gene-level expression between two groups of cells chosen by the user. The cell groups can be chosen from user-provided metadata factors such as sample, tissue or cluster, or UI selections on the UMAP or phylogeny. Users can select cells on the UMAP using a box select or lasso select. Users can select cells on the phylogeny using a lasso select, or by choosing the parent internal node of the cells they wish to include. Users can compare metadata groups or selections to the remaining unselected cells, or to other groups or selections. The log fold change, AUC and FDR-adjusted p value are reported for each gene in the dataset.

### Analysis of the lung cancer data

The tumor phylogeny for CP004 was reconstructed using the Cassiopeia-Hybrid (Jones *et al*., 2020) algorithm from processed single-cell lineage tracing data, as described in the original study (Quinn *et al*., 2021). Cells present in both the expression matrix and the lineage for CP004 were used; otherwise, cells were pruned from the lineage using the *ape* R package (Paradis and Schliep, 2019) or removed from the expression matrix. All unifurcations (i.e., nodes containing exactly one child) were collapsed using the *collapse*.*singles* function in *ape*. Before analysis with *PhyloVision*, the cells of expression matrix of raw UMI counts were library-size-normalized to the median number of UMIs in CP004. Informative genes were found using the *filterGenesFano* function in VISION with default parameters and passed to *PhyloVision* via the *projection_genes* parameter. Signatures were downloaded from the MSigDB database (https://www.gsea-msigdb.org/gsea/msigdb/) and the Hallmark, C2, and C5 (BP) collections were used for analysis. Meta data corresponding to each cell was downloaded from NCBI GEO, series GSE161363 and dataset GSM4905334. *PhyloVision* was run with default parameters, except for setting *num_neighbors = 30* and *projection_methods = c(“UMAP”, “tSNE30”)*. After *PhyloVision* analysis, *Hotspot* was invoked using the *runHotspot* command with the following parameters: *model = “normal”, tree=TRUE, min_gene_threshold=70, n_neighbors = 30, number_top_genes = 1000*.

## SUPPLEMENTARY FIGURES AND LEGENDS

**Figure S1:**
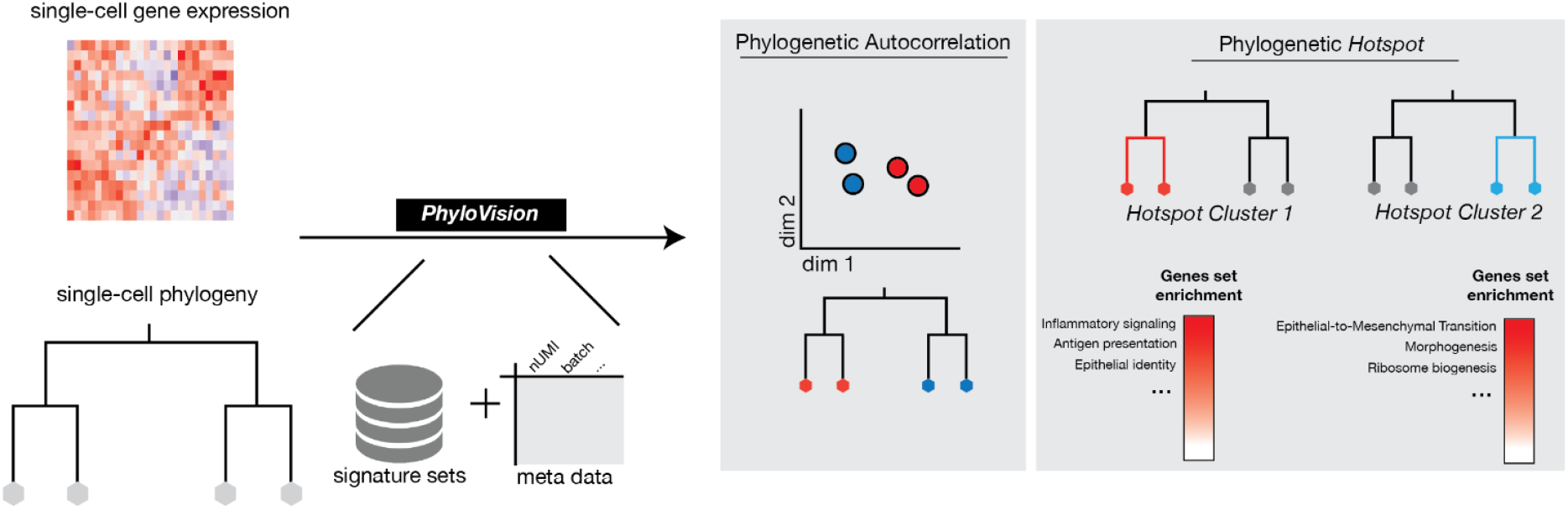
The *PhyloVision* analysis pipeline. A simplified, schematic representation of the *PhyloVision* pipeline. *PhyloVision* takes as input a gene expression matrix, a phylogeny, gene signature sets, and optionally metadata associated with each cell. Signature scores are computed for each cell in the dataset and evaluated with phylogenetic autocorrelation (see Methods). Upon user specification, *PhyloVision* performs *Hotspot* gene module identification using the phylogeny as a latent space. Modules can be interpreted by assessing the enrichment score between signature gene sets and module gene sets.

**Figure S2:**
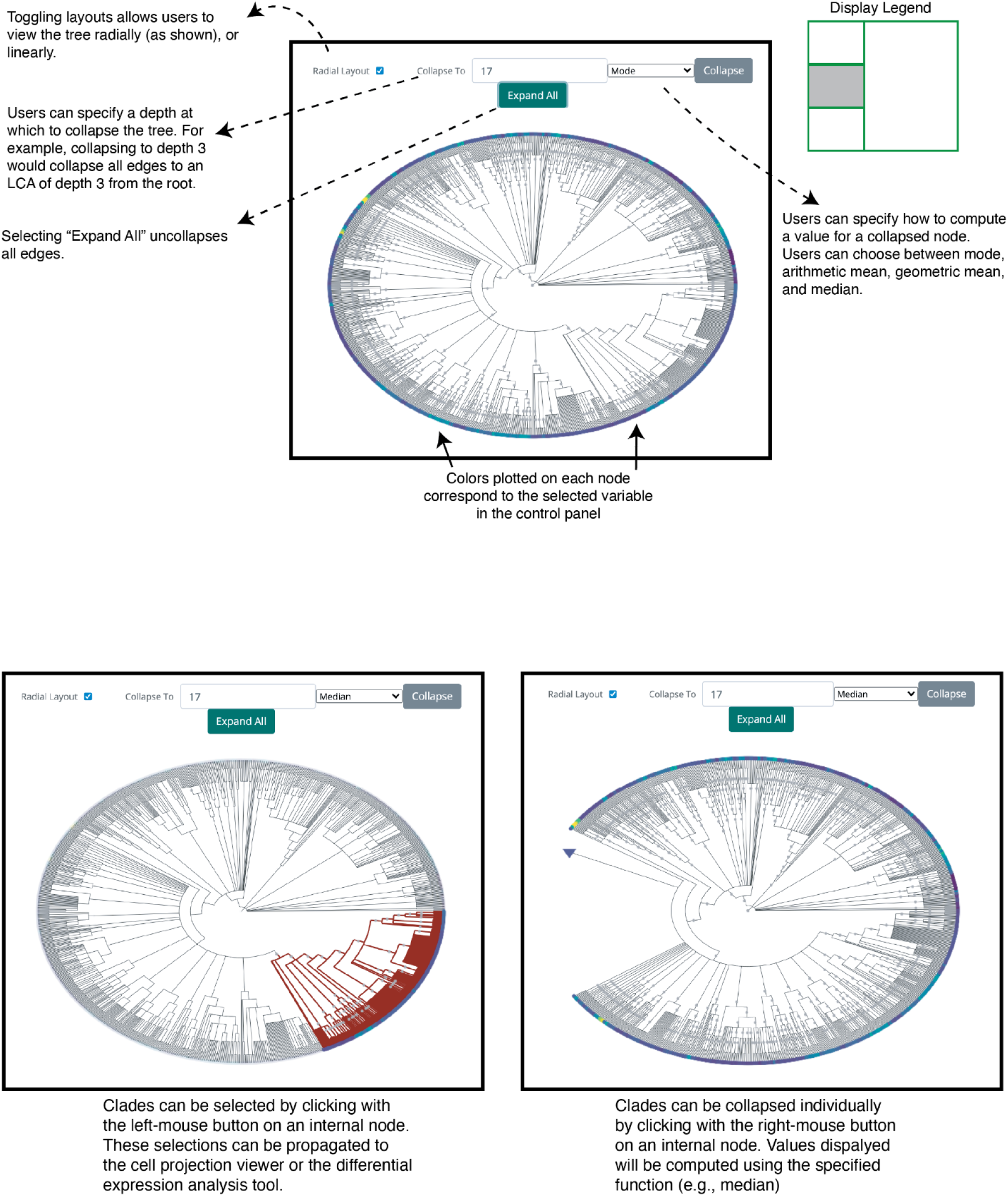
Elements of the PhyloPlot interface. A tutorial over the features of the PhyloPlot.js interface used in the *PhyloVision* web-based report. Screenshots indicate how to interact with the object, including how to select, collapse, and expand clades.

**Figure S3:**
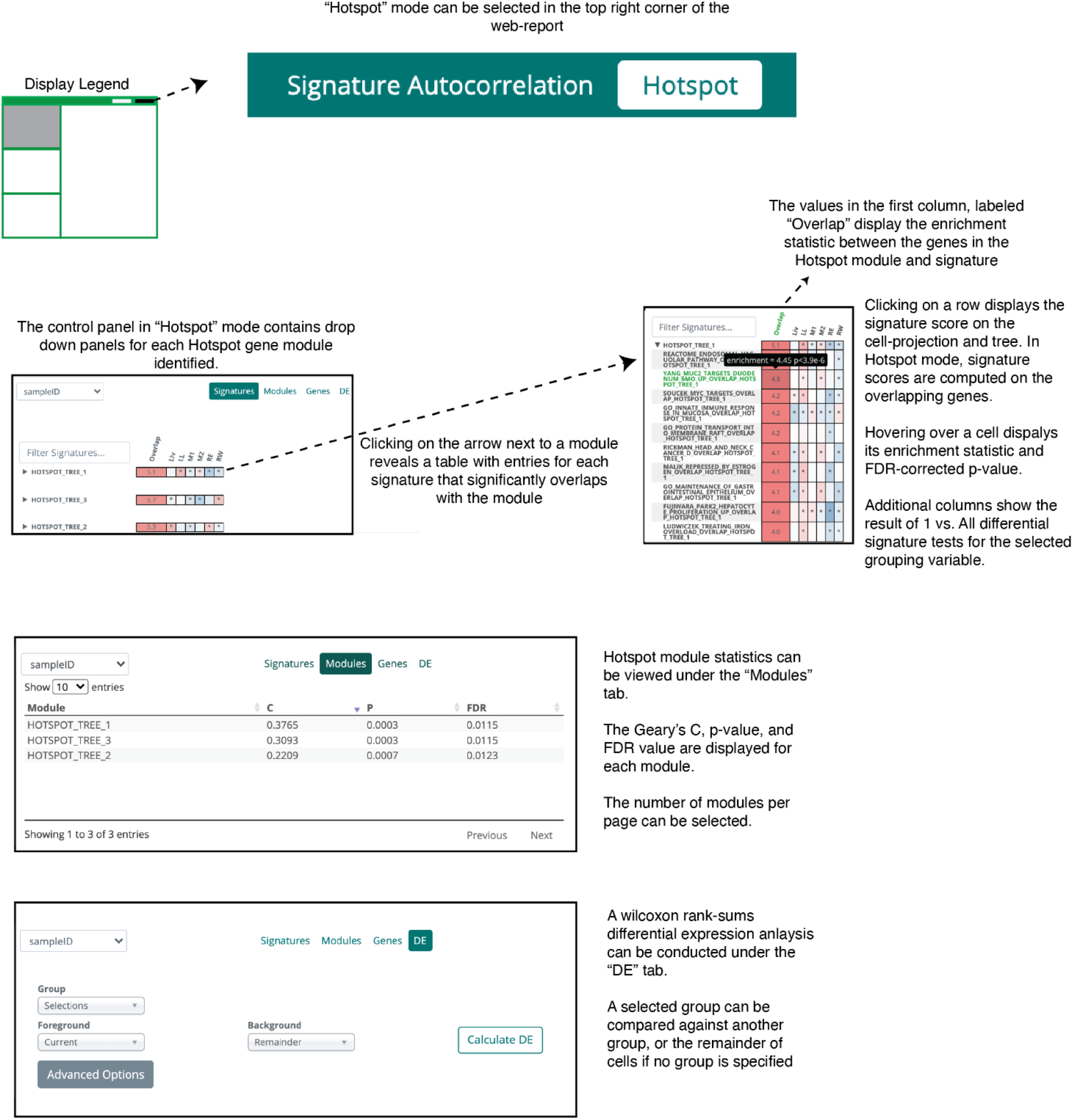
Elements of the *Hotspot* mode control panel. Users can toggle between signature autocorrelation and *Hotspot* modes in the top-right corner of the web-based report. In *Hotspot* mode, each module is presented in a collapsable table. Elements in the table are gene signatures that significantly overlap with the *Hotspot* module. Users can also view the *Hotspot* summary statistics with the “Modules” tab, where the Geary’s C, p-values, and FDR of each module are displayed.

**Figure S4:**
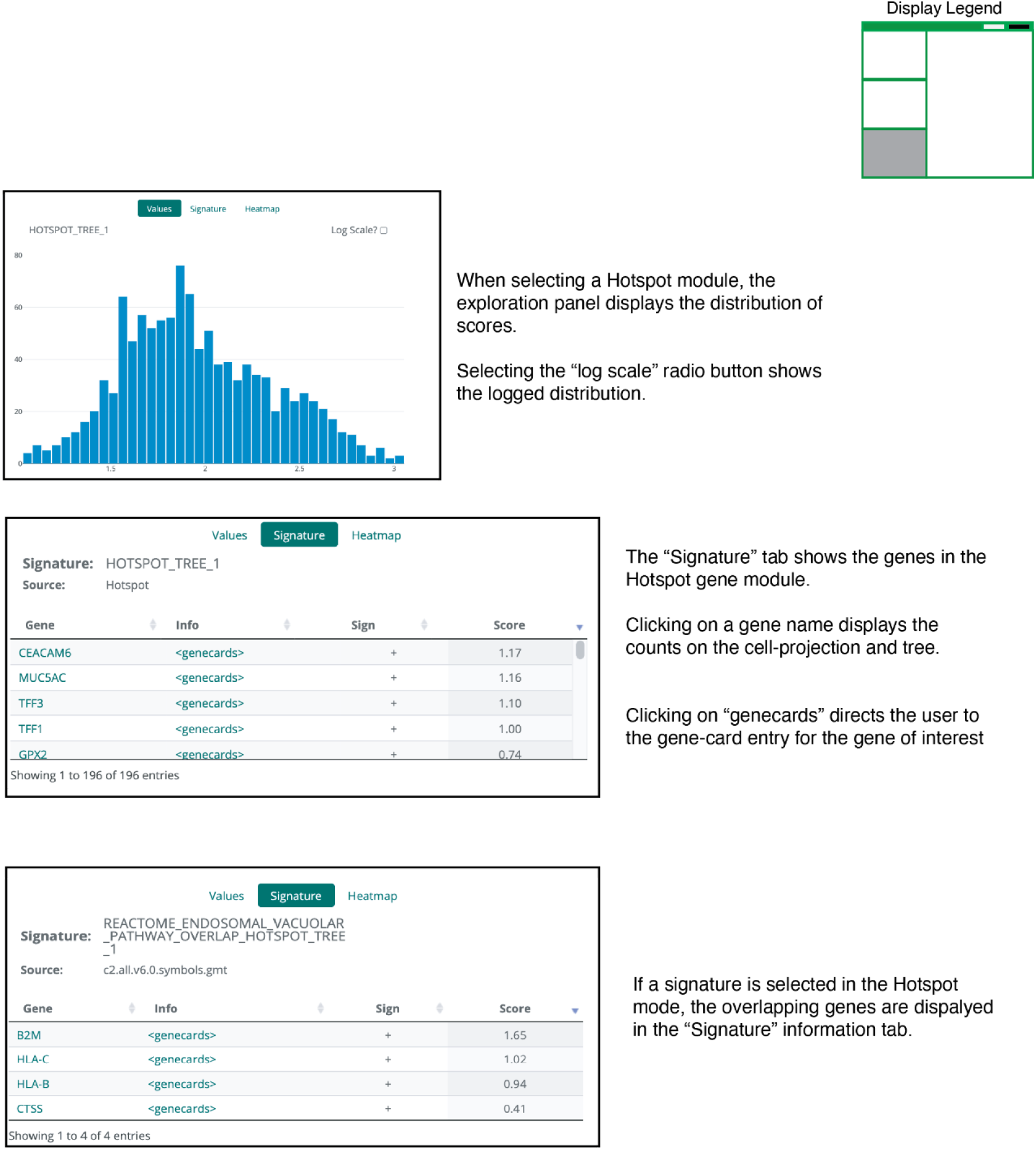
Elements of the *Hotspot* mode signature exploration panel. Users can view the distribution of scores in the *Hotspot* mode on both normal and log scales. Scores are the *VISION-*computed score for the signature consisting of genes overlapping between *Hotspot* module and gene signature. If a signature is selected, the distribution displayed is the signature score of only the genes overlapping between the *Hotspot* module and the gene signature. Users can also view the genes in a *Hotspot* module, or in the overlap between a gene signature and *Hotspot* module.

## Notes

### Competing Interest Statement

The authors have declared no competing interest.

